# High-resolution structural and functional deep brain imaging using adaptive optics three-photon microscopy

**DOI:** 10.1101/2021.01.12.426323

**Authors:** Lina Streich, Juan Boffi, Ling Wang, Khaleel Alhalaseh, Matteo Barbieri, Ronja Rehm, Senthilkumar Deivasigamani, Cornelius Gross, Amit Agarwal, Robert Prevedel

**Affiliations:** Cell Biology and Biophysics Unit, European Molecular Biology Laboratory, Heidelberg, Germany; The Chica and Heinz Schaller Research Group, Institute for Anatomy and Cell Biology, Heidelberg University, Heidelberg, Germany.; Interdisciplinary Center for Neurosciences, Heidelberg University, Heidelberg, Germany.; Epigenetics and Neurobiology Unit, European Molecular Biology Laboratory, Monterotondo, Italy.; Developmental Biology Unit, European Molecular Biology Laboratory, Heidelberg, Germany.; Molecular Medicine Partnership Unit (MMPU), European Molecular Biology Laboratory, Heidelberg, Germany.; Collaboration for joint Ph.D.degree between EMBL and Heidelberg University, Faculty of Biosciences

## Abstract

Multi-photon microscopy has become a powerful tool to visualize the morphology and function of neural cells and circuits in the intact mammalian brain. Yet, tissue scattering, optical aberrations, and motion artifacts degrade the achievable image quality with depth. Here we developed a minimally invasive intravital imaging methodology by combining three-photon excitation, indirect adaptive optics correction, and active electrocardiogram gating to achieve near-diffraction limited resolution up to a depth of 1.2mm in the mouse brain. We demonstrate near-diffraction-limited imaging of deep cortical and sub-cortical dendrites and spines as well as of calcium transients in deep-layer astrocytes *in vivo*.

---

In the scattering tissues such as the mammalian brain, two-photon excitation microscopy (2PM) is the gold standard for recording cellular structure and function in non-invasive and physiologically relevant conditions *in vivo* [1,2]. However, the maximum penetration depth of two-photon microscopes is fundamentally limited by the on-set of out-of-focus fluorescence near the surface with increasing excitation power, which for the mammalian brain prevents imaging beyond ~1mm [3]. Recently, imaging approaches based on three-photon excitation microscopy (3PM) have shown potential for deep imaging beyond 1mm on a cellular scale, due to a significantly increased signal-to-background ratio at depth and longer wavelength excitation [4–7]. Like in 2PM, however, with increasing imaging depths optical aberrations due to tissue heterogeneities and refractive index mismatches as well as subtle motion artefacts due to the animal’s heartbeat effectively degrade image resolution and result in a loss of sub-cellular details. This has so far prohibited, without using highly invasive methods such as GRIN lens implantation [8] or cortical aspiration [9], to resolve fine neural processes, synapses, and sub-cellular Ca^2+^ transients in deep cortical and subcortical areas of the mouse brain *in vivo*. While optical aberrations can be measured and compensated by utilizing adaptive optics (AO) methods [10,11], previous implementations were predominantly based on 2PM and thus limited in their effective imaging depth to below 800µm in the mouse [12–15]. Alternative approaches based on so-called wavefront shaping have the potential to image even further into highly scattering media, but limited field-of-views (FOVs) of only tens of micrometer and/or fast decorrelation times have prohibited useful applications in realistic intravital conditions [16–18]. A further challenge in in-vivo deep brain imaging is the fact that at large tissue depths heart pulsation leads to intra-frame motion artefacts which prevent frame averaging to enhance signal-to-noise ratio (SNR), a standard technique that is essential to reliably resolve small structures such as dendrites and individual spines. To address the above shortcomings, here we developed a minimally invasive intravital imaging methodology based on 3PM, indirect adaptive optics (AO) correction and active electrocardiogram (ECG) gating to achieve near-diffraction limited resolution up to a depth of 1.2mm in the mouse brain. This enabled us to resolve individual synapses down to ~900µm in the cortex, and fine dendritic processes in the hippocampus at >1mm depth, both of which would be invisible without AO and ECG-gating. Furthermore, our non-invasive approach achieved the first in vivo functional characterization of fibrous astrocytes in the white matter and to resolve Ca^2+^ transients in individual microdomains.

Our intravital imaging method is based on a home-built three-photon (3P) laser scanning microscope optimized for 1300nm excitation and the use of broad bandwidth, low repetition-rate lasers (**Fig. 1a, Methods** and **Supplementary Fig. 1**). The 3P fluorescence signal was optimized by ensuring short duration laser pulses (<50fs) and efficient power delivery to the focal plane within the sample. Signal optimization in 3PM is especially important for in-vivo imaging as it has been demonstrated that laser induced tissue heating can lead to physiological changes and long-term tissue damage [7,19]. We first validated the deep-tissue capability of our 3P microscope by imaging neuronal structure and third-harmonic generation (THG) contrast along an entire cortical column of a Thy1-EGFP-M mouse through a chronic glass-window (**Fig. 1b** and **Supplementary Video 1**). We observed high signal-to-background ratios (SBRs), even at deep, sub-cortical layers below the white matter, while ensuring physiologically low (0.5-22mW) average laser power and focal energies (<2nJ) below previously established damage thresholds [7] (**Supplementary Fig. 2**). Despite the high SBR as well as numerical aperture (NA) and thus spatial resolution of our 3P system, however, fine neuronal processes such as dendritic spines are only reliably resolved down to ~500µm, and completely disappear below the highly scattering corpus callosum. This is partly due to physiological brain motion resulting from respiratory and cardiac activities. These brain motion artifacts progressively degrade theoretically achievable in vivo imaging resolution with increasing depth. When coupled with relatively slow frame rates (~Hz) in 3PM, due to the use of low-repetition rate lasers, brain motion results in intra-frame imaging artefacts which are difficult to correct during post-processing. These distortions lead to image blurring and prevents frame-averaging to increase image SBR, resulting in reduced resolution and contrast. Therefore, they present an inherent challenge for in-vivo high-resolution intravital microscopy. To address this, we implemented a prospective image gated acquisition scheme based on a field-programmable gate array (FPGA), which enables active synchronization of the scanners to the cardiac cycle in real-time so that active scanning is paused during peaks of electrocardiogram (ECG) recording (**Methods**). We found that ECG-gating substantially reduced intra-frame motion artefact down to the hippocampus (>1mm depth), and now individual spines could be resolved across the entire cortex down to Layer VI. Furthermore, temporal sequences of frames acquired with ECG gating were more correlated to one another compared to non-gated frames, indicating a higher temporal image stability (**Fig. 1c, Supplementary Fig. 3** and **Supplementary Video 2**). Compared to previous work that used ECG-triggered full frame acquisition [20], our approach is independent of chosen acquisition parameters, does not require additional post-processing, and can readily be applied to large FOV scanning.

**Fig. 1:**
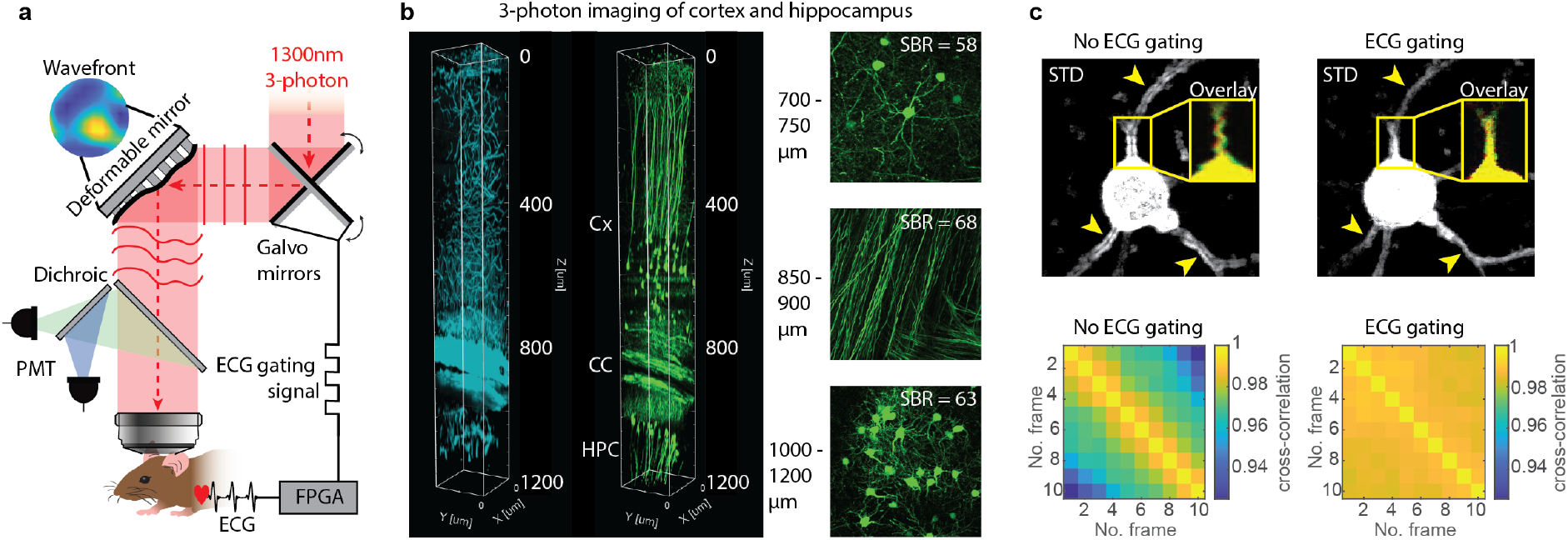
Schematic principle of ECG-gated adaptive optics three-photon microscopy. **(a)** Illustration of motion corrected adaptive optics three-photon microscope. Aberration correction is performed via a modal-based indirect wavefront sensing approach. Intravital motion artefacts are reduced with a real-time ECG gated image acquisition scheme that synchronizes the Galvo scanners to the cardiac cycle of the mouse. **(b)** Three-photon microscopy at 1300nm excitation wavelength in EGFP-Thy1(M) mouse visual cortex and hippocampus. (Left) 3D reconstruction of three-photon image stack of third harmonic signal (cyan) and GFP-labeled neurons (green). (Right) Maximum intensity projection images at various depths in (top) cortex (Cx), (middle) corpus callosum (CC) and (bottom) CA1 region of the hippocampus (HPC). SBR at different depth are displayed in the respective images. **(c)** Comparison of intra-frame motion artefacts for ECG-gated and non-gated image acquisition at 701μm depth. Standard deviation projection images (STD) of consecutively acquired frames with (top, right) and without (top, left) ECG gating. Arrows indicate high frame-to-frame variability resulting in artefacts without ECG gating. Yellow box shows overlay of two consecutively acquired frames (red and green). (Bottom) Pairwise 2D-cross correlation between individual frames without (bottom, left) and with ECG (bottom, right) synchronization, indicating reduced frame-to-frame variability with ECG-gating. Representative datasets obtained from (n=4) mice.

Degradation of the excitation point spread function (PSF) due to optical aberrations is particularly problematic in 3PM, as the fluorescence depends non-linearly on the focal intensity [4]. Furthermore, sub-micrometer structures such as dendrites and spines are smaller than the axial extent of the PSF, and thus become challenging to resolve. Optical aberrations further increase the axial spread of PSF, which leads to overall low fluorescence signal from fine structures. Previous work that utilized 3PM for deep cortical and sub-cortical imaging focused on cellular (functional) investigations that did not require sub-cellular details [5,7,21]. In order to restore near-diffraction limited resolution in deep brain regions, we incorporated adaptive optics to our intravital ECG-gated 3PM. We chose indirect, modal-based AO [22] as these approaches are most robust in low-signal and high-noise conditions [23] and are arguably best suited to push the depth limits in deep brain imaging, especially when combined with higher-order multiphoton microscopy [24]. We developed a deformable mirror (DM-) and modal-based AO optimization scheme with automatic shift correction and integrated it into the hardware control of our 3PM. Then we validated the AO performance by correcting artificial aberrations in ex-vivo brain slices (**Methods** and **Supplementary Fig. 4,5**). Next, we applied our method to *in vivo* 3P imaging of neuronal structures throughout the cerebral cortex and hippocampus of Thy-1 EGFP-M mice (**Fig. 2a,b** and **Supplementary Fig. 6-8** and **Supplementary Videos 3,4**). We observed significant improvements in image quality and the spatial resolution due to AO correction. We directly measured up to 4-fold improvement in effective axial resolution and about 6-fold enhancements of fluorescence signals, as evidenced by axial intensity line plots and spectral power map analysis of lateral resolution (**Fig. 2c,d** and **Supplementary Fig. 7**). In addition, we note that the reported improvements in signal and resolution (*AO full*) are solely due to correction of tissue-induced aberration, since values are compared to best performance of the microscope, including aberrations due to sample preparation (*AO system*). With our 3P-AO scope, we were able to recover near-diffraction limited performance at all imaging depth, and in particular to reliably resolve individual synapses down to ~900µm in the cortex (**Fig. 2a** and **Supplementary Videos 3,4**) and fine dendritic processes in the hippocampus at >1mm depth (**Fig. 2b, Supplementary Fig. 3g** and **Supplementary Fig. 8**), which were invisible without AO and ECG-gating. As a further demonstration, we applied our method to a thinned-skull preparation, which is optically challenging due to significant bone-induced aberrations that normally restrict high-resolution imaging to the first hundred micrometers [25]. We observed similar improvements in image quality (**Supplementary Fig. 9**), as described in recent work utilizing AO and 2PM [26]. A full diffraction limited resolution could not be restored, however, likely due to the presence of higher-order aberrations that were not included in the correction scheme.

**Fig. 2:**
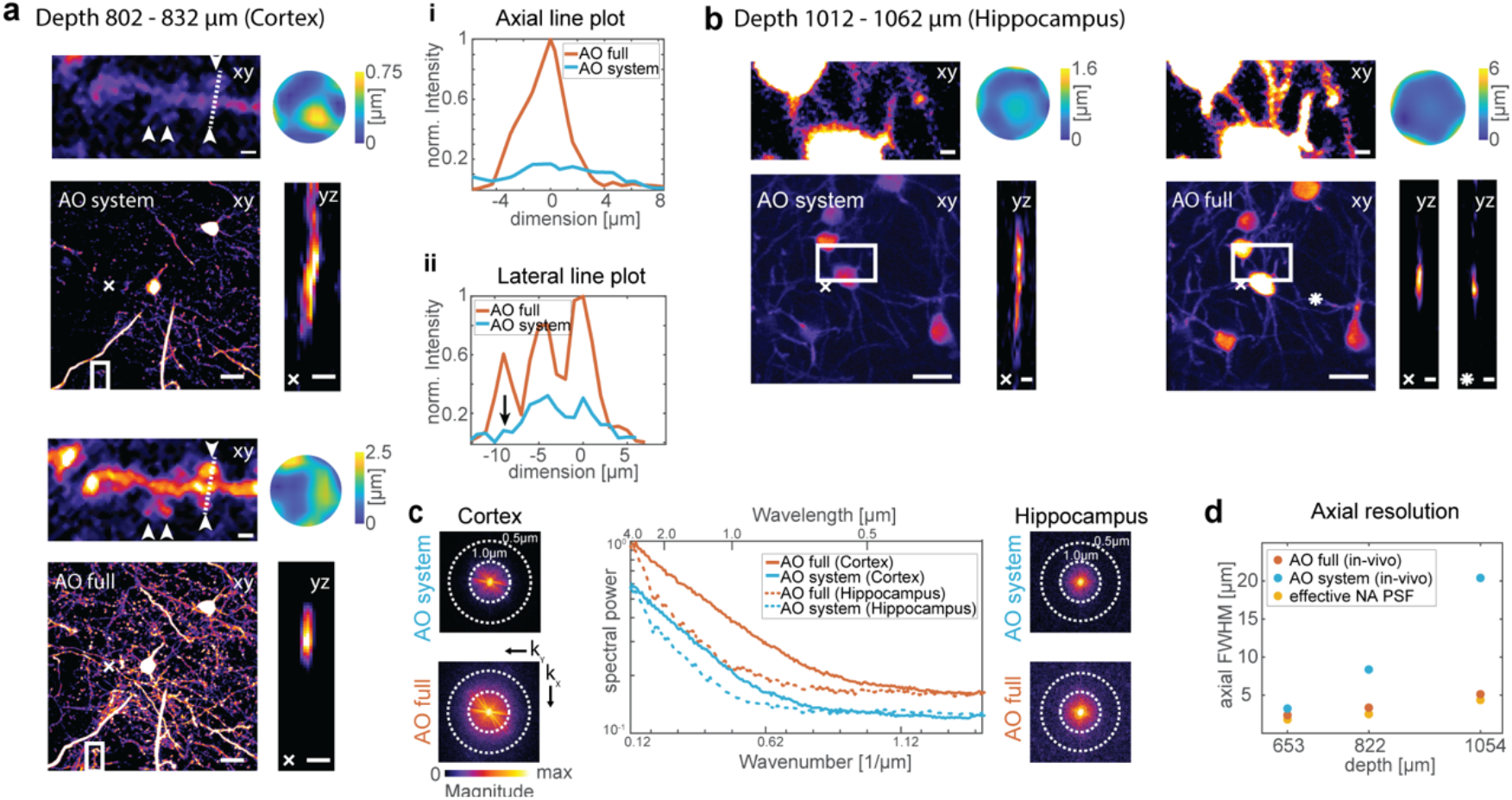
AO enables high-resolution 3P deep brain imaging. Indirect modal-based adaptive optics correction and ECG-gated three-photon microscopy at 1300nm. Representative images showing dendrites and spines in **(a)** layer VI and **(b)** hippocampus of the in-vivo mouse brain (EGFP-Thy1(M)) through a cranial window. Images were recorded with two different conditions: *AO system*: wavefront correction of system aberrations; *AO full*: wavefront correction of system and brain tissue aberrations. Maximum intensity projection images with system and full aberration correction at **(a)** layer VI in cortex as well as **(b)** hippocampus (scale bar 20μm). White boxes indicate magnified view of spines which clearly become visible with full adaptive optics correction. White cross indicates orthogonal view (yz) along dendrite. Respective wavefronts for aberration correction are displayed in the top corners. **(a, i)** Lateral (xy) and **(a, ii)** axial (yz) intensity profile along spines (white dotted line) and across dendrite (white cross), respectively, in cortical layer VI (scale bar 2μm). **(c)** Lateral resolution analysis of AO correction improvement. Spectral power map as a function of spatial frequency (wavenumber) at (left) layer VI of cortex and (right) hippocampus (shown in (b)). (Middle) average radial profile of spectral power maps for maximum intensity images. **(d)** Axial resolution analysis. Effective PSF of microscope inferred from beads in solution and compared to in-vivo measurements performed on dendrites, with system and full adaptive optics at depth 653μm (cortex), 822μm (cortex), and 1054μm (hippocampus) – also see **Supplementary Figure 6,7**. The effective illumination NA was reduced at larger depth to increase objective transmission, yielding slightly decreased resolution. Representative datasets obtained from several imaging sessions in (n=4) mice.

Next, we explored the capability of our 3P-AO scope for an even more challenging application of multi-photon microscopy, namely in studying the physiology of glial cells which are notorious for their low fluorescence in their fine-processes and high photosensitivity [27]. Similar to neurons, astrocytes, the most abundant glial cell-type in the brain, constitute a diverse population of cells with morphological and functional heterogeneity across different brain areas [28]. Two most prominent astrocyte populations, *protoplasmic astrocytes* (PA) and *fibrous astrocytes* (FA) reside in grey- and white matter areas respectively (**Fig. 3a**). While PAs have been extensively investigated non-invasively *in vivo* in the upper cortical layers utilizing common 2P methods [29], little is known about the Ca^2+^ signals of PAs and FAs located in deeper brain areas. The main hurdle in studying FAs has been their location among highly scattering myelinated fibre tracts deep within the brain (>800µm) and the lack of genetic tool-box to target these cells. Moreover, astrocytes are highly amenable to photoactivation, and prolonged exposure to high laser power can severely affect their physiology [27,29–32]. To further demonstrate our technical advancements, we used a AAVs based viral approach to express genetically encoded Ca2+ sensor GCaMP6f specifically in astrocytes, and utilized our 3P-AO microscope to in-vivo record Ca^2+^-transients in FAs in the corpus callosum (**Fig. 3b** and **Supplementary Video 5**) and deep layers (layer V-VI) PAs (**Fig. 3c** and **Supplementary Figure 10**). Here, the use of AO was crucial to achieve high signal and SBR even under low laser excitation power and enabled the detection of more active microdomains compared to without aberration correction (**Fig. 3d** and **Supplementary Videos 6-8**). We note that the overall time of AO optimization was fast (<500ms per independent Zernike mode, see **Methods**) compared to Ca^2+^-dynamics lasting for several seconds in the astrocyte soma, this allowed us to use indirect AO on structures with functionally varying signal levels. We resolved Ca^2+^ transients in individual microdomains, which are linked to synaptic activity, and yielded high signal-to-noise ratios even at low power levels. To our knowledge, this is the first recording of *in vivo* sub-cellular Ca^2+^ transients in individual FAs in subcortical white matter (**Fig. 3b**). Our approach enabled the exciting discovery that, although FAs reside in the white matter and do not tend to form ‘regular’ synaptic contacts as seen in PAs, they still exhibit microdomain-like Ca^2+^ transients.

**Fig. 3:**
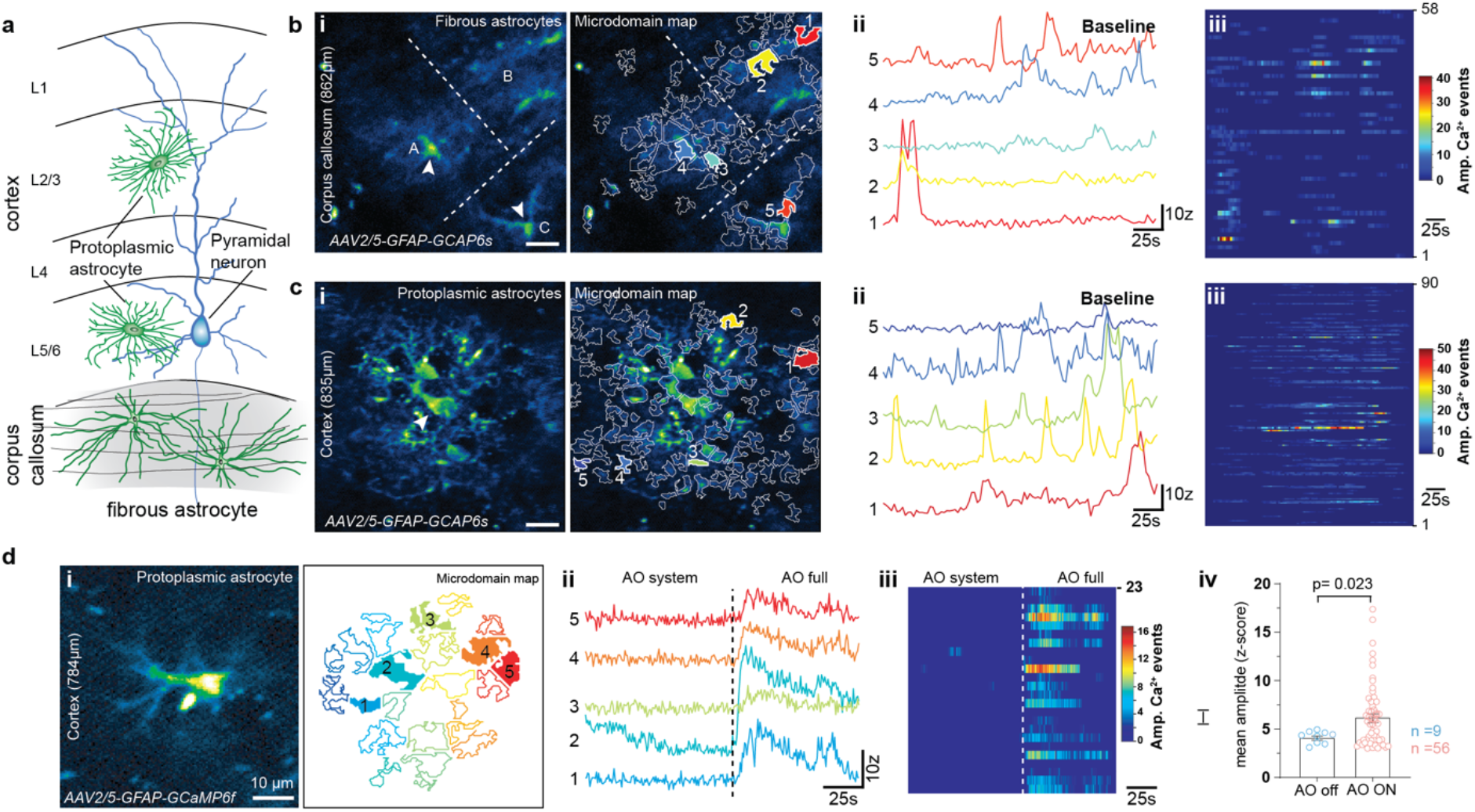
3P-AO enables structural and functional imaging of astrocytes *in vivo*. **(a)** Illustration of protoplasmic and fibrous astrocytes which reside in the grey and white matter, respectively. **(b)** White matter astrocytic Ca^2+^-imaging. (**i**) Median intensity time-series projection image (pseudocolored, left) of three fibrous astrocytes (A, B, C) in the corpus callosum (at depth of 862µm). Arrowheads show soma of astrocytes. Map of all active microdomains at baseline overlaid on median intensity projected image of astrocyte (right). (**ii**) Intensity versus time traces for five microdomains (corresponding to colors in (i, right), showing characteristics of Ca^2+^ transients. (**iii**) Raster plots displaying duration and intensity of Ca^2+^ transients of all microdomains. **(c)** Grey matter astrocytic Ca^2+^-imaging. (**i**) Median intensity time-series projection image (pseudocolored, left) of 2 protoplasmic astrocytes in the layer 6 of the visual cortex (at depth of 835µm). Map of all active microdomains at baseline overlaid on median intensity projected image of astrocyte (right). (**ii**) Intensity versus time traces for five microdomains (corresponding to colors in (i), right), showing characteristics of Ca^2+^ transients. (**iii**) Raster plots displaying duration and intensity of Ca^2+^ transients of all microdomains. **(d)** AO-enhanced astrocytic Ca^2+^-imaging. (**i**) Median intensity time-series projection image (pseudocolored, left) of protoplasmic astrocyte in the cortex (at 784 µm depth). Map of active microdomains active at baseline (right). (**ii**) Intensity versus time traces for five microdomains (corresponding to colors in (i), showing characteristics of Ca^2+^ transients without AO (AO off) and full AO (AO on) correction. (**iii**) Raster plots displaying duration and intensity of Ca^2+^ transients without AO and full AO correction. (**iv**) Graph showing mean amplitude (Z score/SNR) for microdomain Ca^2+^ transients without AO and full AO correction; p-value based on non-parametric Kolmogorov-Smirnov unpaired t-test. Representative datasets shown from (n=3) mice, and (n=5) L5/6 protoplasmic cells (c), (n=4) fibrous cells (b), and (n=3) cells with adaptive optics (d).

To summarize, by combining three-photon microscopy with active aberration correction and ECG-gating, we advanced deep-tissue microscopy for near diffraction-limited *in-vivo* brain imaging in mice. Our demonstrations show the capability of our method to capture neuronal morphology and astrocyte Ca^2+^-signals at a resolution not previously reported at these depths. In line with recent reports [13,33], we achieved clear resolution enhancements utilizing AO throughout the cortex and the hippocampus (at >1mm depth), while maintaining low-power levels necessary for physiological recordings of subcellular and photosensitive processes [7]. Although the indirect AO approach used in this work is in general slower than direct wavefront sensing methods, our independent modal-based optimization is sufficiently fast to be used on samples with slowly varying signal levels, including Ca^2+^-signals in astrocytes. This is a unique advantage of our method compared to other approaches [17,18,33] that require fluorescence levels to stay constant over much longer time-frames. Furthermore, indirect AO is not sensitive to the large chromatic differences between excitation and signal emission, and not depth-limited by wavelength of the (shorter) fluorescence light, and allowed AO optimization to be performed at relatively low excitation power levels (see **Methods**). Our methods allow investigation of new biological questions, such as layer specific synaptic remodeling which previously was restricted to the first three layers of the brain, required removal of overlying brain tissue [9] or insertion of invasive, optical probes [34]. Furthermore, we showed the capability of our method to uncover the distinct physiology of astrocytes across different cortical layers and of a previously understudied population of sub-cortical white matter astrocytes. Apart from our chosen demonstrations, we believe that our intravital 3P-AO technology can be easily extended to other tissues and across various model organisms in order to investigate cellular structure and physiology (and pathophysiology) with minimal invasiveness. In the future, wavefront shaping approaches [17], which can correct for both scattering and aberrations simultaneously will have great potential when combined with 3PM to maintain diffraction-limited performance at even larger depths.

## Acknowledgements

We would like to thank the EMBL Heidelberg mechanical and electronic workshop, especially C. Kieser, as well as the animal facilities for help and support. We further thank Jacopo Antonello, Jingyu Wang and Martin Booth from Oxford University for sharing the design and software of the calibration interferometer used in this work. We thank Hannah Sonntag and Rolf Sprengel from Max Planck Institute for Medical Research, Heidelberg, for providing the rAAV1/2 particles expressing gfaABC1D-GCaMP6f. This work was supported by the European Molecular Biology Laboratory (L.S., L.W., M.B., R.R., C.G. and R.P.). J.B. and S.D. were supported by EC Marie Curie COFUND EIPOD3 interdisciplinary postdoctoral fellowships (S.D. was co-hosted by C.G. and R.P.). A.A. is supported by the Chica and Heinz Schaller Research Foundation and grants A09N/SFB1158 and P8/FOR2289 from the Deutsche Forschungsgemeinschaft.

## Author contributions

L.S. built the imaging system, performed experiments, analyzed data and wrote indirect AO and DM calibration software. J.B. performed cranial surgeries and virus injections. L.W. contributed with optical simulation, mechanical design and to DM calibration software. K.A. analyzed astrocyte Ca^2+^-imaging data under supervision of A.A. M.B. developed ECG-gating software together with L.W. R.R. performed skull thinning preparations under supervision of J.B.. S.D. and C.G. contributed to project design and preliminary experiments. A.A. conceptualized and designed astrocyte imaging experiments. R.P. conceived, designed and supervised the project. R.P., L.S. and A.A. wrote the manuscript with input from all authors.

## Methods

### Excitation source

The 3-photon excitation source was a wavelength-tunable non-collinear optical parametric amplifier (NOPA, Spectra Physics) pumped by a regenerative amplifier (Spirit, Spectra Physics) operated at 1300nm and 400kHz. At this wavelength the laser system delivered an output power of 470mW with a pulse width of 32fs. To compensate for group delay dispersion (GDD) of the microscope optical components we implemented a homebuilt single-prism pulse compressor [35], which consisted of a N-SF11, 40mm prism (PS855, Thorlabs) and two gold roof mirrors (HRS1015-M01, Thorlabs). With this pulse compressor ~45fs pulses (deconvolved with a sech^2^ pulse shape) could be recovered at the objective focal plane, as measured by an autocorrelator (Carpe, APE GmbH). The excitation power was modulated with a Pockels cell (360-40 LTA, ConOptics).

### Custom-built adaptive optics multi-photon microscope

A schematic of the microscope set-up is shown in **Supplementary Fig. 1**. For adaptive optics correction the deformable mirror (for details see below) was conjugated to the back focal plane (BFP) of a high-NA objective (NA 1.05, 25x, XLPLN25XWMP2, Olympus) via a pair of relay lenses (for DM-Multi-3.5 (Boston Micromachines), L1, L2: 47-380, Edmund Optics; for DM97-15 (Alpao), L1: AC508-250-C, L2: AC254-75-C, Thorlabs) to galvanometers (GVS002, Thorlabs), which were conjugated to each other with a relay system of two paired lenses (lens L3: 47-380, Edmund Optics, L4: LE4950, Thorlabs). The galvometers, scan lens, tube lens and objective BFP were conjugated in ‘4f’-configuration. The scan lens was composed of 2” lenses (AC508-100-C, Thorlabs) in Ploessl configuration and a broad-bandwidth tube lens (TL200-2P2, Thorlabs). The microscope head and detection unit were based on a Movable Objective Microscope (MOM, Sutter Instrument Company), and three-dimensional translation of the objective was achieved with an electronic micromanipulator translation stage (MP-285). The fluorescent signal was collected by the same objective, separated from the excitation path with a long-pass dichroic beam splitter (D1: FF76-875, AHF) and filtered with a near-infrared laser blocking filter (FF01-940/SP, Semrock). The THS and fluorescent signal were separated with a dichroic beam splitter (D2: T480lpxr, AHF), cleaned with optical filters (F1: FF01-432/36-30-D, Semrock; F2: ET525/70m-2p, AHF), respectively, and detected by a photomultiplier tube (Hamamatsu, H7422-40), amplified and low-pass filtered by a transimpedance amplifier (3.5MHz bandwidth, 10^5^x gain, Femto DHCPA-200). The amplified voltage signals were digitized at a sampling rate of 80MHz by an A/D conversion module comprising a digitizer adapter (NI-5734, National Instruments) and an FPGA card (PXI-7961R, National Instruments). ScanImage 2016a (Vidrio Technologies, [36]) running on MATLAB 2016a (MathWorks) was used to control image acquisition and was custom modified to enable synchronization of the image acquisition to the laser pulse repetition rate, integrated by the FPGA to realize a scheme in which each voxel is illuminated by a single or constant multiple of laser pulses. Synchronization to the laser repetition rate also allowed the introduction of a signal sampling window (125ns was found optimal) that ignores noisy samples where no signal was present, thus improving the signal-to-noise ratio of the collected data.

### Deformable mirror calibration

For active wavefront (WF) modulation two different continuous membrane deformable mirrors (DMs) were used: a DM-Multi-3.5 (Boston Micromachines) and the DM97-15 (Alpao) with 140 and 97 actuators, respectively. Calibration and generation of a Zernike-to-Command control matrix (Z2C) was performed with a homebuilt Michelson-interferometer to calibrate the Boston-DM while the Alpao-DM was inhouse calibrated with a homebuilt Shack-Hartmann wavefront sensor (SHWS), consisting of a sCMOS camera (C11440-22CU, ORCA Flash4.0, Hamamatsu) with a 20×20 microlens array (18-00197, SUSS MicroOptics) and custom written control software (additionally the Alpao-DM was pre-calibrated in the factory). Continuous membrane DMs have non-linear response behavior which results in the Z2C matrix being linear only around the empirically chosen amplitude range used for calibration. To ensure a reliable operation even outside this range, we characterized the effect of Zernike modes on the axial displacement of the PSF. Axial shifts of the image plane are problematic in modal-based adaptive optics correction as this can lead to changes of the metric value due to content change of the image rather due to change of the excitation PSF and hence can lead to sample dependent biases [37]. This may corrupt the optimization procedure and prevent convergence to an optimal solution. We observed that independent of the calibration procedure, both DMs showed cross-talk between spherical modes and defocus. Therefore, a look-up table was generated for the Z2C matrix after calibration that characterized the respective focal shift for the 1st, 2nd and 3rd order spherical mode. For moderate mode amplitudes (~60% of maximum RMS value) mode coupling was linear, therefore the focus shift for spherical modes could be compensated by applying the opposite focal shift directly to the DM using the Zernike defocus mode (**Supplementary Fig. 4)**. However, due to non-linear mode coupling this focus shift correction failed for large amplitudes. Hence, for large amplitude aberrations the axial focal shift of spherical modes was compensated by moving the microscope translation stage along the axial dimension. This approach was integrated into our inhouse custom-written indirect AO software (described below) and the axial translation was based on the generated focal shifts look-up table.

### Modal-based indirect wavefront optimization

For wavefront (WF) optimization, a user-defined number (N) of Zernike modes were corrected independently and the DM WF updated sequentially after optimization of each mode. For each mode 9 amplitude changes were applied during the first optimization round and 5 amplitude modulations during subsequent iterations. As a quality metric for the WF optimization we chose the mean intensity of user-defined structures of interest such as cell soma, as this showed the most robust performance in low SBR conditions. Furthermore, we found cell soma to be less prone to sample dependent biases [37] and because of their brightness allowed using exceptionally low-power during the AO routine. During optimization, the structures of interest were segmented and only the segmented image was analyzed. Segmentation was achieved by intensity thresholding the median filtered (3×3pixel) image. Zernike modes were modulated in the following order: 1st spherical, 2nd coma, 3rd astigmatism followed by other modes. This modulation scheme was chosen because spherical and coma aberrations are common and known to have large amplitudes in in-vivo cranial window imaging as they result from refractive index mismatches between layers and tilted surfaces, respectively [38]. In general, two to three AO iterations were performed before convergence of the optimization metric was observed. Hence, in total 9N+(i-1)*5N measurements were performed (i, number of iterations, N number of Zernike modes) to find the optimal WF. Normally, tip/tilt and defocus modes were excluded from the correction scheme, however at large depths (> 1000µm) the defocus mode was included into the correction scheme to reposition the focal plane onto the object of interest after every iteration as vital functions of the animal could lead to translational brain motion and hence sample dependent biases. This can lead to an elongation of the PSF towards the brightest structure if it is not centered at the initial focal plane [37].

### Typical parameters for 3P-AO optimization

For indirect modal-based WF optimization the focal plane was positioned in the center of the brightest structure in the region-of-interest (ROI). In principle, structures of different size may be used for wavefront optimization (as validated on fluorescent beads of size 2-10µm - data not shown). For in-vivo experiments, we chose neurons (Thy1-GFP mice) or astrocyte (expressing GCaMP6f) somata for indirect AO optimization. Typically, a 40μmx40μm FOV was scanned with a 64×64 pixel resolution, resulting in a ~25Hz acquisition rate during optimization. Independent mode optimization was essential for aberration measurement on dynamic astrocyte calcium signals. With an independent mode optimization scheme the fluorescent signal only needs to be stable over 5-9 amplitude modulations. This data could be acquired within ~250-450ms, which is at least one order of magnitude faster than somatic astrocyte GCaMP dynamics [about 8.0s, [39]]. For cranial window preparations, 21-36 Zernike modes, excluding tip/tilt mode, were used for aberration correction and 2-3 iterations were performed. Hence, a total of 266-646 measurements were required for aberration measurement which resulted in an overall fluorescence acquisition time of 15-34s at 25Hz frame rate. Acquisition parameters for AO optimization and imaging are summarized in **Supplementary Table 2**. To minimize brain heating and photobleaching during AO optimization, the excitation power was reduced substantially, approx. by 50%, so that only a dim signal of the large somata was visible. Utilizing previously determined parameters for the light attenuation l_z_ at 1300nm in brain tissue (l_z_=270µm, see Ref. [6]) the estimated focal energy E_f_ was ~1nJ (E_f_ = E_o_*exp(-z/l_z_), z: depth from surface) during WF optimization which is within the established physiological pulse energy regime [7]. Although we employed physiological power and pulse energy regimes, aberration measurements were performed on a different astrocyte within the proximity of the astrocyte of interest used for functional recordings, in order to ensure physiological responses during the imaging period. Furthermore, to prevent experimental biases and activation of astrocytes during recording the order of AO correction (first AO off followed by AO on, or vice versa) was chosen randomly (**Supplementary Video 6-8**). For structural imaging in Thy-1-EGFP mice, the WF correction and imaging was performed on the same cell somata. Furthermore, to increase the objective transmission at large depth the effective illumination NA was reduced (NA~0.8), yielding slightly decreased resolution (**Fig. 2d**).

To minimize aberrations before AO optimization, the correction collar of the objective was adjusted to compensate for the refractive index mismatch of the cranial window. Furthermore, the cranial window was aligned orthogonal to the optical axis employing an objective-like alignment tool which enabled alignment within <1deg accuracy. The alignment tool was based on overlaying the reflection from the cranial window to a reference reflection of a surface perpendicular to the optical axis at >2m distance.

### ECG-gating

#### Software development

ScanImage was custom modified to acquire images gated in real-time to the physiological parameters of the mouse, such as breath and heart rate, which were monitored by a dedicated physiological monitoring system (ST2 75-1500, Harvard Apparatus). The ECG-signal triggered a TTL pause gate (generation of the gate is described below) which was used to modify the system’s laser clock which was fed into the FPGA through an auxiliary I/O connector block (SCB-19, National Instruments). When the state of the pause gate was set to high, this effectively halted the image acquisition task and the galvometers stopped at their current position. During this period, the Pockels cell was synchronously turned off via an external circuit based AND gate in order to prevent extensive light exposure of the sample. Scanning continued once the pause gate was reset to low.

#### Hardware development

The ECG-triggered TTL pause gate was developed on an FPGA System-on-Chip platform (ZYBO Z7 ZYNQ-7010, Digilent) using Verilog programming language on Vivado 2018. First, the analog ECG signal was scaled to the FPGA input range (0-1V), using an operational amplifier to reduce the peak-to-peak amplitude of the signal and centering it at 0.5V. Next, the ECG signal R-wave peaks (the local highest amplitude in the ECG signals) of the cardiac cycle was taken as the base point to generate the pause gate. The status of the TTL pause gate was set to low within the time interval [t_p_-Δt_pre_, t_p_+Δt_post_], where R wave peak occurred at time t_p_ (**Supplementary Fig. 3a**). The parameters Δt_pre_ and Δt_post_ were manually adjustable in accordance with suppression of the motion artifacts in the images. In our experiments, Δt_pre_ and Δt_post_ were set to 30% and 20% of the ECG-cycle period, resulting in an imaging duty cycle of 50%. In practice, it is hard to predict the exact occurrence time of the next R wave peak. Therefore, we continuously analyzed the last 10 heart beat cycles to predict the next R wave peak in real-time. Furthermore, we developed an auto-tuning threshold algorithm which adapted the threshold levels to changing DC levels of the ECG signal.

### Animals and ethics statement

This work followed the European Communities Council Directive (2010/63/EU) to minimize animal pain and discomfort. All procedures described in this manuscript were approved by EMBL’s committee for animal welfare and institutional animal care and use (IACUC), under protocol number RP170001. Experiments were performed on C57Bl6/j or homozygous Thy1-EGFP-M (Jax No: 007788) transgenic mice from the EMBL Heidelberg core colonies. During the course of the study mice were housed in groups of 1-5 in makrolon type 2L or 3H cages, in ventilated racks at room temperature and 50% humidity while kept at a 12/12 hr light dark cycle. Food and water was available *ad libitum*.

### Stereotaxic viral vector delivery and cranial window implantation

For the the astrocyte specific expression of GCaMP6f, the gfaABC1D-cyto-GCaMP6f expression construct (Addgene: Cat#52925) was sub-cloned into the AAV vector. rAAV serotypes 1 and 2 viral particles were generated as described previously [40] and purified by AVB Sepharose affinity chromatography [41] after titration with real-time PCR (each titer 1.0 – 6.0E12 viral genomes/ml; TaqMan Assay; Applied Biosystems). For stereotaxic injection of rAAVs and the cranial window implantation, mice were anesthetized with isoflurane vapor mixed with O_2_ (5% for induction and 1-1.5% for maintenance). Anesthetized mice were subcutaneously injected with 1% xylocain (AstraZeneca) under the scalp as preincisional local anesthesia and placed in a stereotaxic frame (David Kopf Instruments, model 963). The skin and periosteum over the dorsal cranium was removed with fine forceps and scissors to expose the bone. A 4mm diameter circular craniectomy was made over the visual cortex using a dental drill (Microtorque, Harvard Apparatus), centered at 2.5mm posterior and 2.5mm lateral to Bregma. Close care was taken to preserve the integrity of the dura and avoid bleeding. rAAV injections were performed at the center of the craniectomy using glass pipettes lowered to depths of 400, 800 and 1200 μm, using a syringe at a rate of ~4ul/hr. ~300nl were injected per spot. After injection, a round 4mm coverslip (~170μm thick, disinfected with 70% ethanol) was placed over the craniectomy with a drop of saline between the glass and the dura. The craniectomy and cranial window were sealed using acrylic dental cement (Hager Werken Cyano Fast and Paladur acrylic powder), and a customized headbar was cemented to the skull for head fixation under the microscope. The skin wound around the surgical area was also closed with dental acrylic. After surgery mice received pain relief (Metacam, Boehringer Ingelheim) and antibiotic (Baytril, Bayer) injections (subcutaneous, 0.1 and 0.5 mg/ml respectively, dosed 10μl/g body weight). Mice were single housed after cranial window implantation and had a recovery period of at least 3 weeks before further experiments, for GCaMP6f expression and to resolve the inflammation associated with this surgery [42].

### Thinned skull and brain slice preparation

Skull thinning followed the same general steps as the cranial window implantation procedure described above. Once the skull was exposed, a round ~3mm area centered at 2.5mm posterior and 2.5mm lateral to Bregma was thinned using a dental drill. The thinned area was deepened a few microns beyond the trabecular part of the bone and the thinned surface was leveled out with polishing dental drill bits, leaving ~25-30 μm of remaining bone tissue. The thickness of the tinned area was estimated by visualization of blood vessels through the progressively more transparent thinned skull.

For brain slice preparation, Thy1-EGFP-M mice were fully anesthetized with isoflurane vapor, 5% in O2 and transcardially perfused with saline followed by 4% paraformaldehide (PFA). Perfused brains were dissected and post fixed overnight at 4°C in 4% PFA. Fixed brains were washed and stored in saline at 4°C until slicing. Free floating 300 μm thick coronal sections of the fixed brains were produced in a vibratome (Leica VT1200S) and mounted on glycerol for imaging.

### In vivo deep brain imaging

For in vivo imaging experiments, animals were head-fixed using a customized headbar and complement holder and the cranial window was aligned orthogonal to the optical axis using the method described above. For structural neuronal imaging, Thy1-EGFP mice were anesthetized with isoflurane (2% in oxygen, Harvard Apparatus) while for functional astrocyte imaging mice were kept under low anesthesia (isoflurane 1-1.5% in oxygen). Mice were positioned on a small animal physiological monitoring system (ST2 75-1500, Harvard Apparatus), which allowed to maintain animal body temperature at 37.5°C and to monitor physiological parameters, ECG signal and breath rate. During experiments eyes were covered with eye ointment. Acquisition parameters for structural neuronal imaging and functional astrocyte Ca^2+^ imaging are summarized in **Supplementary Tables 1 and 2**. The data shown in the Figures is representative of several imaging sessions. In general, the imaging sessions were repeated between 2 to 11 times each on 8 separate mice, all of which produced comparable data quality. The numbers of mice are indicated in the Figure captions.

### Image post-processing and alignment

All image data shown represent either raw or 3×3 median filtered images, as stated in the respective captions and summarized in **Supplementary Table 1**. To improve visibility of dim structures before AO correction a scaling factor was applied to images (also indicated in the figures). Slow image drifts due to translational brain motion were corrected by template matching individual image frames. Structural neuronal images from Thy1-eGFP mice were registered by riding body transformation with StackReg [43] (ImageJ) while astrocyte time-series were registered by translation with TurboReg (ImageJ).

### Automated extraction and analysis of astrocyte Ca^2+^ signals

Astrocytes Ca^2+^ signals were analyzed using a custom-built MATLAB based machine learning algorithm called CaSCaDe (Calcium Signal Classification and Decoding, for details see Ref. [27]). Since CaSCaDe was designed for the analysis of 2-Photon microscopy data, here we incorporated the following changes to optimize the code for analyzing 3P microscopy data: a 3D bandpass filter of 11×11×11 pixels was applied to the time series image, the sum-intensity projected image was binarized using a threshold value of I_bg_, the image was smoothed with a 31×31 pixels^2^ Gaussian filter and a standard deviation of 3 pixels. To generate a microdomain map, a watershed segmentation with more than two regional maxima was applied, and microdomains with an area of less than 5×5 pixels^2^ were considered as noise and discarded. A Ca^2+^ event was considered positive if it had an intensity value of 3 (a.u.) and it spanned more than 4 frames. A support vector machine (SVM) was trained to further refine the detection of Ca^2+^ events. 75 features were extracted from raw and two smooth intensity profiles obtained using a Gaussian filter with sizes of 11 and 21 frames separately. SVM training was performed using MATLAB on 4300 fluorescence signals manually annotated as positive or negative.

### Code availability

The indirect adaptive optics routines developed in this work for integration into ScanImage [32] and the FPGA code developed for the ECG-gating are available from the corresponding author upon reasonable request.

### Data availability

The datasets generated and/or analysed during this study are available from the corresponding author upon reasonable request.

