## Supplementary Information for "High-resolution structural and functional deep brain imaging using adaptive optics three-photon microscopy"

#### Supplementary Videos

**Supplementary Video 1: 3D image stack visualization incl THG signal of Thy1-GFP mouse visual cortex and hippocampus down to 1.2mm depth.** 3D reconstruction (Imaris) of 3p image stack of GFP-labeled neurons (green) in the visual cortex and hippocampus and the THS (cyan). Frames were normalized and segmented for visualization. The corpus callosum and hippocampus start around 825µm and 925µm, respectively. Same data as shown in Fig. 1b.

**Supplementary Video 2: Comparison of ECG and non-gated image acquisition.** Images were acquired at 701µm depth at 1Hz frame rate without (left) and with ECG-gating (right). Heartbeat induced intra-frame motion artefacts which appear as wave-like distortions are minimized with ECG-gating.

**Supplementary Video 3: Comparison of system and full AO correction at layer VI.** Images stack recorded from 807-833µm depth with ECG-gated acquisition. Fine structures and spines become visible with full AO (right) correction which cannot be resolved with system (left) correction only.

**Supplementary Video 4: Comparison of system and full AO correction at layer VI.** Images stack recorded from 786-840µm depth with ECG-gated acquisition. Fine structures and spines become clearly visible with full AO (right) correction.

**Supplementary Video 5: 3P-AO imaging of baseline $Ca^{2+}$ transients in protoplasmic astrocyte *in vivo*.** Astrocytes in the layer VI of primary visual cortex (V1) at a depth of 835µm imaged in an adult mouse. Images were acquired at 2 frame/s, and are displayed at 10 frame/s. Pseudocolored time series image sequence (left) and location and number of microdomains inferred by CaSCaDe analysis (right). Image size 120x120µm. Same data as shown in Fig. 3c.

**Supplementary Video 6: AO improves signal and SBR in astrocyte $Ca^{2+}$-imaging.** Astrocytes in the layer VI of primary visual cortex (V1) at a depth of 784µm imaged in an adult mouse. Images were acquired at 1.37Hz, and are displayed at ~10 frame/s. At 200.75s, the AO is switched on. Image size 58x58µm. For visualization purposes 5 frames were averaged and the image was smoothed. Same data as shown in Fig. 3d.

**Supplementary Video 7: AO improves signal and SBR in astrocyte $Ca^{2+}$-imaging.** Astrocytes in the layer VI of primary visual cortex (V1) at a depth of 782µm imaged in an adult mouse. Images were acquired at 1.37Hz, and are displayed at ~10 frame/s. At 109.50s, the AO is switched on. Image size 92x92µm. For visualization purposes 5 frames were averaged and the image was smoothed.

**Supplementary Video 8: AO improves signal and SBR in astrocyte $Ca^{2+}$-imaging.** Astrocytes in the layer VI of primary visual cortex (V1) at a depth of 767µm imaged in an adult mouse. Images were acquired at 1.37Hz, and are displayed at ~10 frame/s. At 186.15s, the AO is switched off. Image size 59x59µm. For visualization purposes 5 frames were averaged and the image was smoothed.

### Supplementary Figures

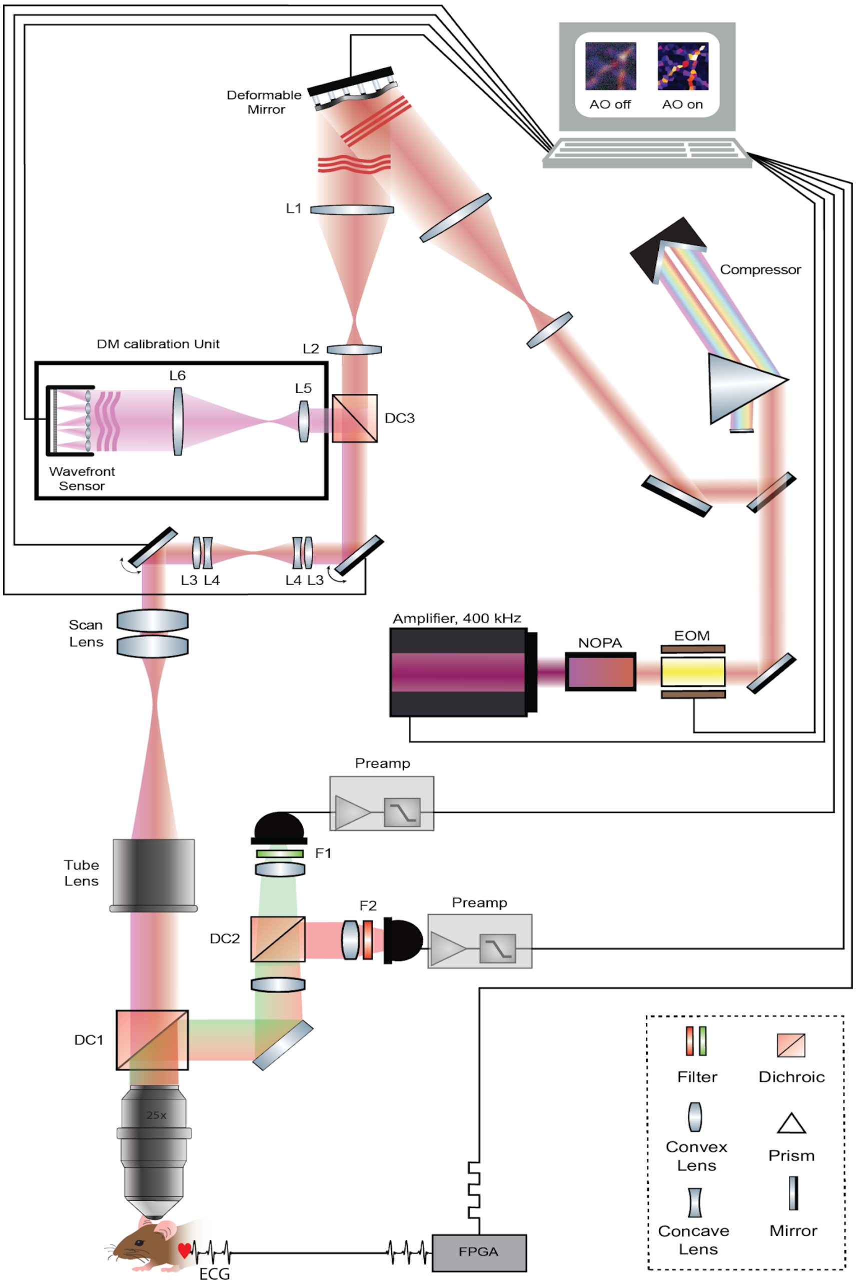


**Supplementary Figure 1: Schematic experimental set-up**. Femto-second laser pulses at 1300nm are generated by a NOPA system pumped by an fiber-based amplified laser. Laser pulse duration is maintained at the sample plane with a custom-build single-prism pulse compressor [1]. The deformable mirror, X and Y galvo mirrors, and the wavefront sensor are conjugated to the objective pupil plane with 4-f relay lens pairs. Fluorescence signals are collected by large aperture optics and detected by large area PMTs. Physiological parameters of the mouse (ECG, breath rate, etc.) are processed in real-time by an FPGA which in turn gates the image acquisition of the microscope. For details see Methods.

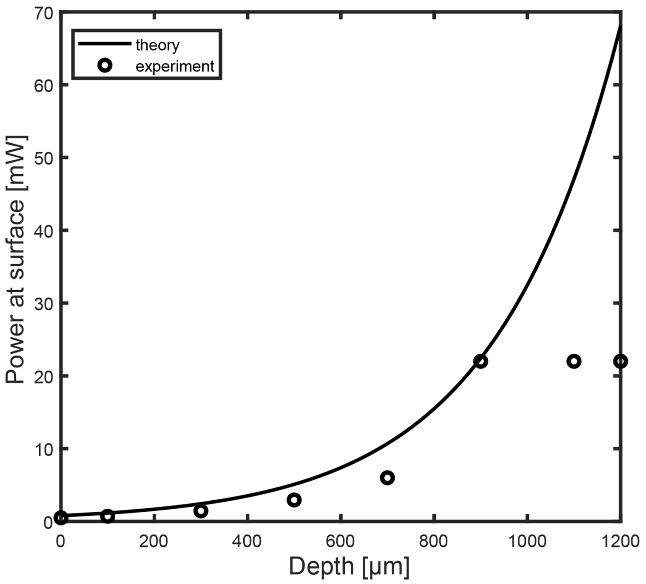


**Supplementary Figure 2: Depth dependent power and focal energy.** Experimental power values used during typical imaging experiments for different depths. Straight line: theoretically determined power $P_0$ at the sample surface for a pulse energy of 2nJ at the focal plane for different depth z. $P(z) = P_0*\exp(-z/l_z)$; attenuation length $l_z$ = 270µm [2].

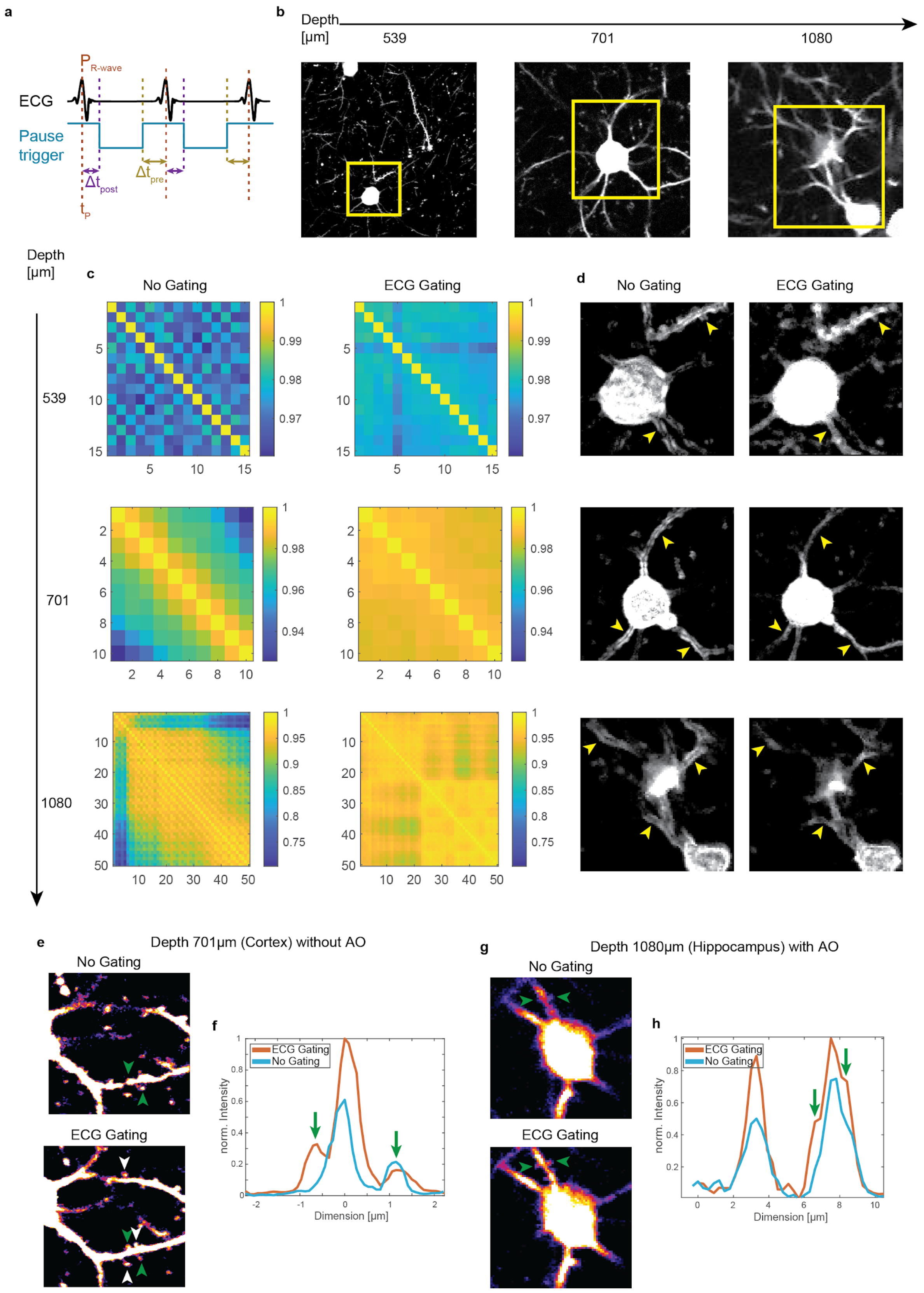


**Supplementary Figure 3**: **ECG-gating reduces intra-frame motion artefacts in in-vivo three-photon microscopy. (a)** Schematic of the trigger signal (cyan) that gates image acquisition (low during acquisition, high during pause time). ECG R-wave peak (brown) at time point $t_p$, capping time $\Delta t_{post}$ (purple), time difference to the predicted next heart beat, $\Delta t_{pre}$ (green). **(b)** Average intensity projection of consecutively acquired frames with ECG-gated image acquisition for different depth and acquisition parameters: depth 57µm, 256x256 pixel, pixel dwell 25µs; depth 536µm, 512x512 pixel, pixel dwell 15µs; depth 701µm, 512x512 pixel, pixel dwell 12.5µs; depth 1080µm, 128x128 pixel, pixel dwell 25µs. **(c)** 2D-cross correlation matrix between pairwise individual frames without (left) and with (right) ECG synchronization. **(d)** Standard deviation projection images (STD) of consecutively acquired image frames with (right) and without (left) ECG gating. Boundaries, indicated by yellow arrows become clearly visible without synchronization while no pronounced boundaries are visible with ECG gating, indicated by yellow arrows. **(e,g)** Averaged three-photon image of **(e)** 15 consecutively acquired frames (FOV = 34µm) at depth 701µm without adaptive optics correction and **(g)** 10 consecutively acquired frames (FOV = 42µm) at depth 1080µm with adaptive optics correction, and corresponding line plots across spines indicated by green arrows in **(f)** and **(g)**, respectively. Spines, indicated by white and green arrows, become clearly visible with cardiac gated acquisition while these fine structures are blurred out without image synchronization.

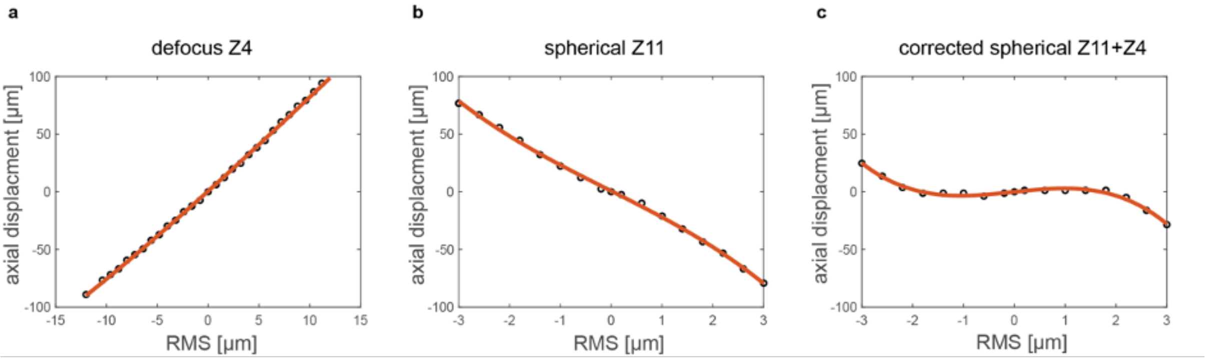


**Supplementary Figure 4**: **Axial focus shift calibration of DM.** Results shown for DM97-15 (Alpao) and measured with 2µm fluorescent beads for different mode amplitudes for Zernike **(a)** defocus mode Z4 and **(b)** first spherical mode Z11. **(c)** The axial focus shift can partially be compensated (for small amplitudes) by combining the defocus and spherical mode.

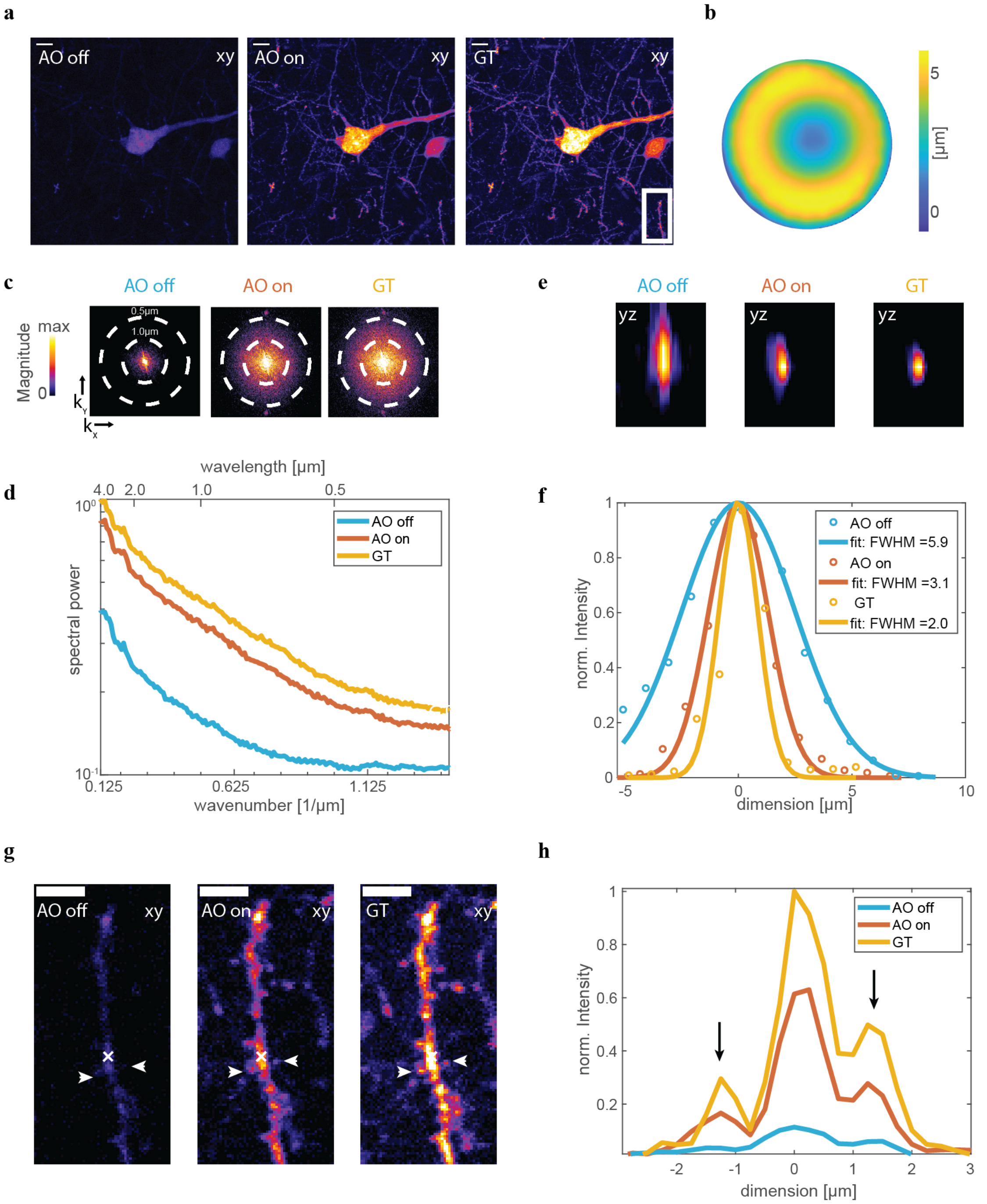


**Supplementary Figure 5: Validation of indirect adaptive optics in ex-vivo Thy1-EGFP(M) brain slices.** Correction of wavefront aberrations introduced by detuning the objective correction collar. Images were recorded with three different conditions. AO off: detuned correction collar and flat DM; AO on: WF correction and detuned correction collar; GT: optimal correction collar with system aberration correction. **(a)** Maximum Intensity projection image for the three different conditions over 20µm depth. **(b)** Corrected wavefront (WF) of Alpao DM. **(c)** Spectral power as a function of spatial frequency (wavenumber) for the images in (a). **(d)** Average radial profile of spectral power maps in c). **(e)** Orthogonal view along selected dendrite indicated in g) by white cross. **(f)** Axial (x,z) intensity profile along dendrite displayed in e) with Gaussian fit to determine the FWHM. **(g)** Magnified views of postsynaptic spines and dendrites corresponding to the boxed region in (a). **(h)** Lateral (x,y) intensity profile along spines indicated in g) by white arrows. Scale bar in a) and g) 5µm. Scale bar in e) 2µm.

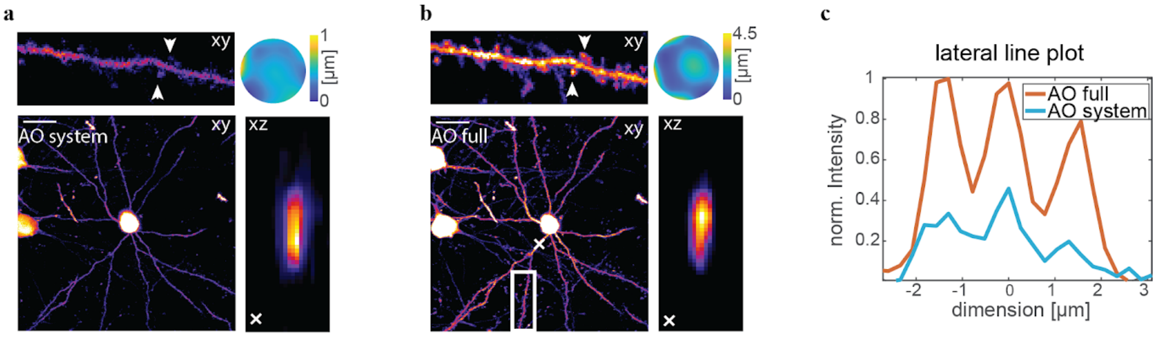


**Supplementary Figure 6: Indirect adaptive optics correction and three-photon microscopy imaging of dendrites and spines in layer V at 653µm depth.** Images were recorded in Thy1-EGFP(M) mouse cortex in-vivo through a cranial window with ECG-gating and two different conditions. AO system: wavefront correction of system aberrations; AO full: wavefront correction of system and brain tissue aberrations. Maximum intensity projection images at 623-653µm depth with **(a)** system and **(b)** with full adaptive optics correction. White boxes indicate magnified view (shown on top) of spines which clearly become visible with full adaptive optics correction. White cross indicates orthogonal view (xz) along dendrite. Respective wavefronts for aberration correction are displayed in the top corners. **(c)** Lateral (xy) intensity profile along spines (white dotted line) (scale bar 20µm).

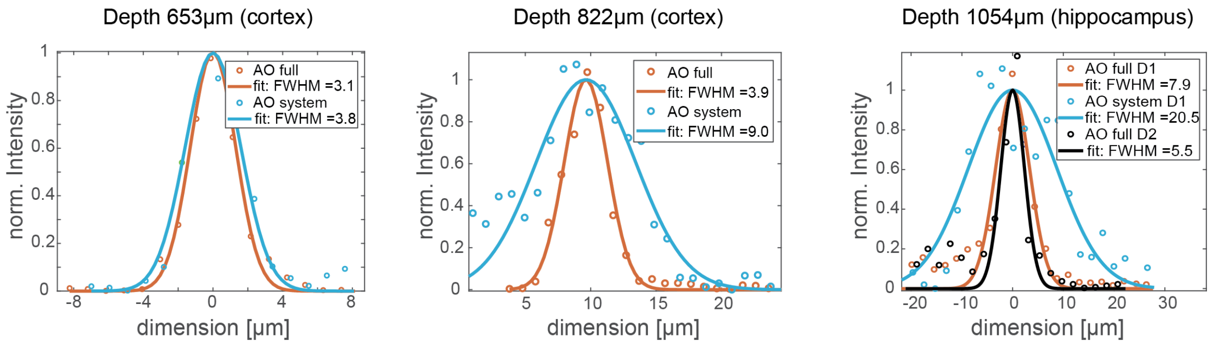


**Supplementary Figure 7: Estimated axial resolution of 3P-AO microscopy at depth in the in vivo-mouse brain with system and full adaptive optics correction.** Axial (xz/yz) intensity profile at depths of 653µm, 822µm and 1054µm (hippocampus) along dendrite shown in Supplementary Figure 6a,b, Figure 2a, and Figure 2b, respectively. Dotted line: experimental data, straight line: Gaussian fit to the data to determine the FWHM which is displayed in the figure legend in micrometer.

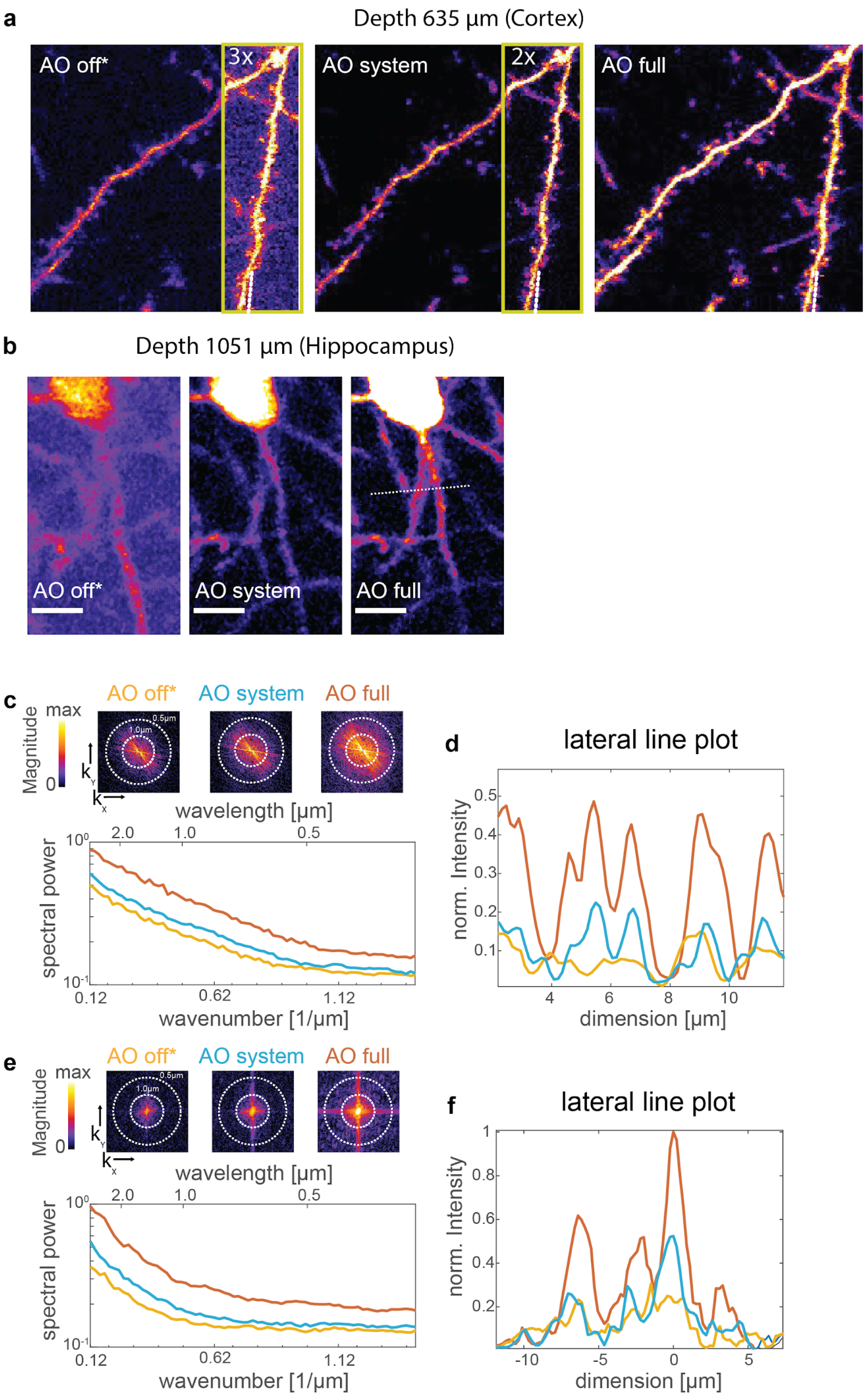


**Supplementary Figure 8: Image quality improvement for in-vivo imaging with ECG-gated acquisition and aberration correction.** Images were recorded in Thy1-EGFP(M) mice with three different conditions. AO off*: no wavefront correction and no acquisition gating; AO system: wavefront correction of system aberrations and cardiac gated image acquisition; AO full: wavefront correction of system and brain tissue aberrations and cardiac gated image acquisition. Maximum intensity projection images for the three different conditions at **(a)** layer V and **(b)** in the hippocampus. FOVs indicated by green box were intensity scaled by stated factor to improve visibility. **(c)** and **(e)** (Top) Spectral power map as a function of spatial frequency (wavenumber) and (bottom) average radial profile of spectral power maps for maximum intensity images and **(d)** and **(f)** lateral intensity plot along dotted line indicated in (a) and (b), respectively.

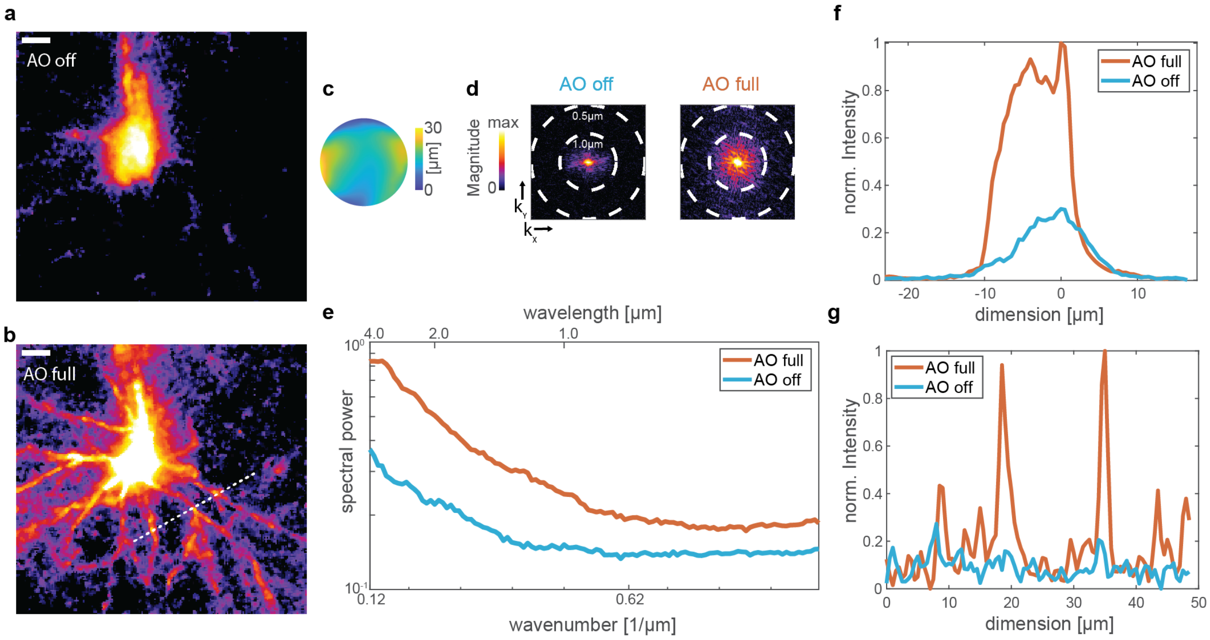


**Supplementary Figure 9: Indirect adaptive optics correction and three-photon microscopy imaging of dendrites and somata through a ~40µm thick thinned skull at 400µm depth below the pia.** Images were recorded in Thy1-EGFP(M) mice with two different conditions: AO off: no aberration correction and AO full aberration correction of Zernike modes up to the 35th mode excluding the tip/tilt and defocus mode. Maximum intensity projection images **(a)** without and **(b)** with aberration correction. **(c)** Corrected wavefront (WF). **(d)** Spectral power map as a function of spatial frequency (wavenumber) of images in (a) and (b). **(e)** Average radial profile of spectral power maps. **(f)** Lateral (x,y) intensity profile along somata and **(g)** along dendrites indicated in (b) by white dotted line. Scale bar in a) 5µm.

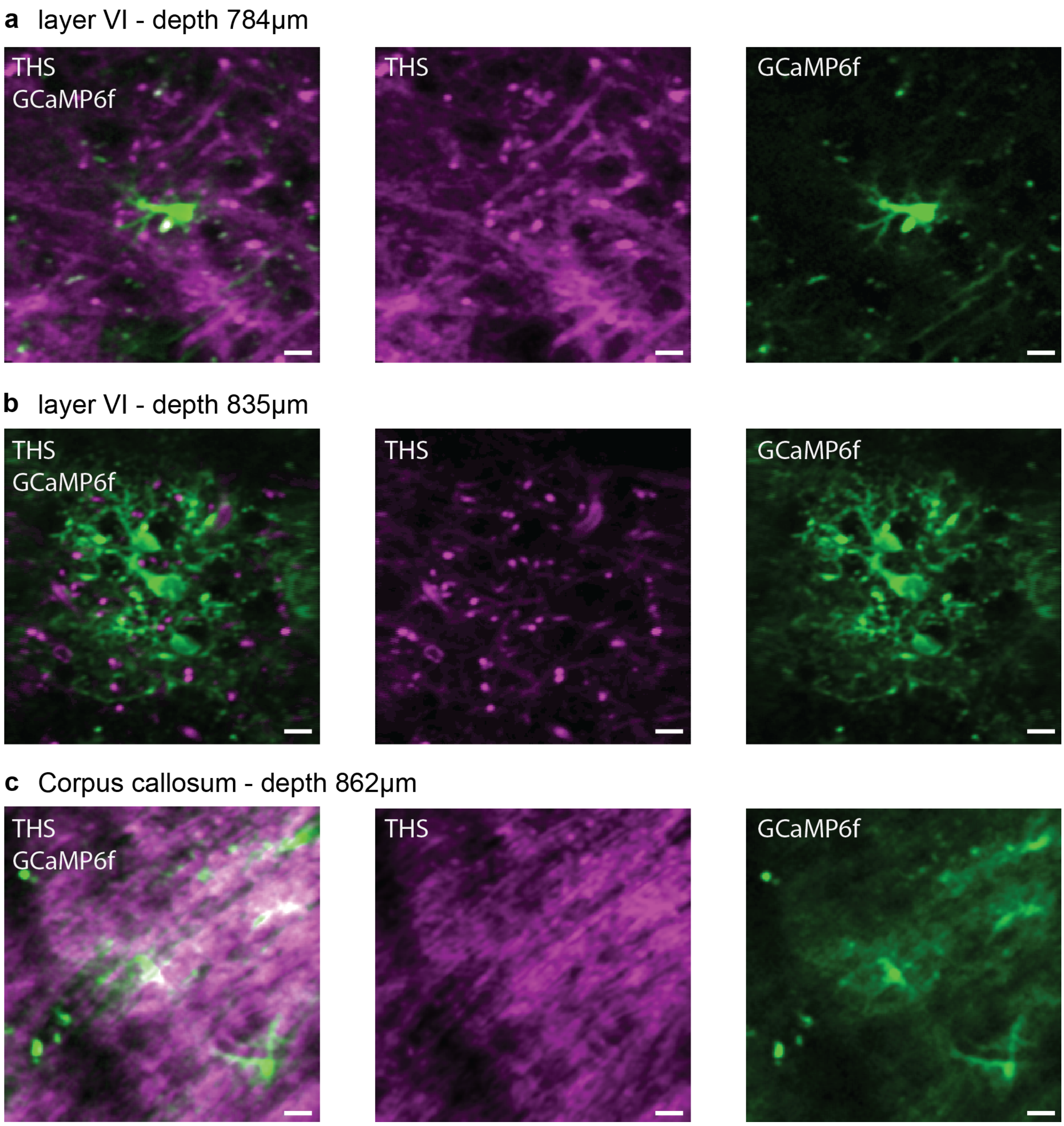


**Supplementary Figure 10: GCaMP6f labeled astrocytes and THS in grey and white matter.** Intrinsic THS generated at the interface of materials with different third-order susceptibility such as blood vessels and the myelinated axons running through the corpus callosum (white matter, c). (a),(b) and (c) correspond to images in Fig. 3d,c and b, respectively.

### Supplementary Tables

Supplementary Table 1: Acquisition parameters for high resolution in-vivo imaging at different depth shown in this work.

| Fig. | Imaging depth (µm) | Power at sample surface (mW) | Image FOV (µm³) | Number of pixels / z-depth increment (µm) | Pixel dwell time (µs) | Number of frames averaged | Frame Rate (Hz) / for time series | Number of frames / for time series | Image postprocessing for visualization |
|---|---|---|---|---|---|---|---|---|---|
| 1b | 0-1200 | 0.5-22 | 215x215x1200 | 512x512/4 | 7.5 | 3 | | | 3x3 median |
| 1c | 701 | - | 50x50 | 512x512 | 12.5 | 1 | 1.09 | 15 | 3x3 median |
| 2a | 822 | 14 | 184x184x30 | 512x512/1.5 | 37.5 | 5 | | | 3x3 median/ Lateral magnification (xy) first scaled by 2, then 3x3 median |
| 2b | 1054 | 30 | 114x114x50 | 512x512/2 | 17.5 | 5 | | | 3x3 median |
| 3b | 862 | 24 | 114x114 | 256x256 | 10 | 1 | 1.37 | 300 | median intensity projection |
| 3c | 835 | 14 | 135x135 | 256x256 | 7.5 | 1 | 1.87 | 400 | median intensity projection |
| 3d | 784 | 9 | 104x104 | 256x256 | 10 | 1 | 1.37 | 500 | median intensity projection |
| **Supp. Fig.** | | | | | | | | | |
| 2b(left) | 539 | 4 | 220x220 | 512x512 | 15 | 1 | 0.23 | 15 | 3x3 median |
| 2b(middle) | Same dataset shown in Fig. 1c | | | | | | | | 3x3 median |
| 2b(right) | 1080 | 22 | 70x70 | 128x128 | 25 | 1 | 2.18 | 50 | 3x3median |
| 2e | Same dataset shown in Fig. 1c | | | | | | | | |
| 2g | 1081 | 22 | 42x42 | 256x256 | 25 | 1 | 0.55 | 10 | raw |
| 4 | Brain slice | - | 130x130x20 | 512x512/2 | - | 2 | | | raw |
| 5 | 653 | 10 | 135x135x30 | 512x512/2 | 25 | 3 | | | raw |
| 7a | 635 | 10 | 30x30x6 | 206x206/2 | 25 | 3 | | | raw |
| 7b | 1050 | 30 | 31x50x10 | 143x229/2 | 17.5 | 5 | | | raw |
| 8 | 400 | 18 | 125x125x30 | 256x256/2 | 10 | 5 | | | 3x3 median |
| **Supp. Video** | | | | | | | | | |
| 1 | Same dataset shown in Fig. 1b | | | | | | | | Imaris |
| 2 | Same dataset shown in Fig. 1c | | | | | | | | smooth |
| 3 | Same dataset shown in Fig. 2a | | | | | | | | smooth |
| 4 | 817 | 9-11 | 164x164x54 | 512x512/2 | 20 | 3 | | | smooth |
| 5 | Same dataset shown in Fig. 3b | | | | | | | | raw |
| 6 | Same dataset shown in Fig. 3d | | | | | | | | smooth/ AVG 5 frames/ scaled by 3 |
| 7 | 782 | 26 | 164x164 | 256x256 | 7.5 | 1 | 1.37 | 300 | smooth/ AVG 5 frames/ scaled by 3 |
| 8 | 767 | 28 | 70x70 | 256x256 | 7.5 | 1 | 1.37 | 500 | smooth/ AVG 5 frames/ scaled by 2 |

Supplementary Table 2: Acquisition parameters for adaptive optics correction at different depth shown in this work.

| Fig. | Imaging depth (μm) | Imaging power at surface (mW) | Frame rate during AO optimization (Hz) | Number of modes corrected | Iteration | Total number of frames |
|---|---|---|---|---|---|---|
| 2a | 822 | 10 | 38 | Z5-Z33 | 3 | 551 |
| 2b | 1054 | 20 | 38 | Z4-Z22 | 2 | 266 |
| 3d | 804 | 9 | 20 | Z4-Z22 | 3 | 361 |
| **Supp. Fig** | | | | | | |
| 4 | Brain slice | - | 38 | Z5-Z22 | 5 | 522 |
| 5 | 653 | 5 | 11 | Z5-Z22 | 3 | 342 |
| 7a | 635 | 5 | 11 | Z5-Z22 | 3 | 342 |
| 7b | 1050 | 20 | 38 | Z4-Z22 | 2 | 266 |
| 8 | 400 | - | 11 | Z4-Z36 | 3 | 627 |
| **Supp. Video** | | | | | | |
| 3 | Same dataset shown in Fig. 2a | | | | | |
| 4 | 817 | 4 | 38 | Z4-Z36 | 3 | 627 |
| 6 | Same dataset shown in Fig. 3d | | | | | |
| 7 | 826 | 20 | 20 | Z4-Z22 | 3 | 361 |
| 8 | 735 | 20 | 20 | Z4-Z22 | 3 | 361 |
